# Chromatin-associated RNA sequencing (ChAR-seq) maps genome-wide RNA-to-DNA contacts

**DOI:** 10.1101/118786

**Authors:** Jason C. Bell, David Jukam, Nicole A. Teran, Viviana I. Risca, Owen K. Smith, Whitney L. Johnson, Jan Skotheim, William J. Greenleaf, Aaron F. Straight

**Author notes:** Correspondence and requests for materials should be addressed to: Jason C. Bell & Aaron F. Straight, &, Beckman Center for Molecular & Genetic Medicine B409, Stanford University, 279 West Campus Drive Stanford, CA 94305.

## Abstract

RNA is a critical component of chromatin in eukaryotes, both as a product of transcription, and as an essential constituent of ribonucleoprotein complexes that regulate both local and global chromatin states. Here we present a proximity ligation and sequencing method called **Ch**romatin-**A**ssociated **R**NA **seq**uencing (ChAR-seq) that maps all RNA-to-DNA contacts across the genome. ChAR-seq provides unbiased, *de novo* identification of targets of chromatin-bound RNAs including nascent transcripts, chromosome-specific dosage compensation ncRNAs, and genome-wide trans-associated RNAs involved in co-transcriptional RNA processing.

## Introduction

Much of the eukaryotic genome is transcribed into non-coding RNA (ncRNA), and several studies have established that a subset of these ncRNAs form ribonucleoprotein complexes that bind and regulate chromatin^1–3^. Some of the most well studied ncRNAs are those involved in dosage compensation, which include *roX1* and *roX2* in *Drosophila* and *Xist* in mammals. In *Drosophila, roX1* and *roX2* are part of the male-specific lethal (MSL) complex that coats the single male X chromosome to induce H4K16 acetylation and increase transcription^4^. In female mammals, *Xist* is expressed from a single locus on the X and coats one of two X chromosomes in order to silence transcription^5^. Other ncRNAs, such as HOTAIR^6,7^ HOTTIP^8^, and enhancer RNAs^9^, have been shown to regulate expression of specific genes by localizing to chromatin and recruiting activating or repressing proteins. Finally, repetitive ncRNA transcripts have roles at chromosomal loci essential in maintaining genomic integrity over many cell divisions, including TERRA at telomeres^10^ and alpha-satellites near centromeres^11^. Despite these well-studied examples, other functional ncRNAs are likely yet to be discovered, the genomic targets of most chromatin-associated ncRNAs are unknown, and the mechanisms by which these ncRNAs regulate chromatin remain largely unexplored.

Genomic methods for studying the localization of specific RNA transcripts include ChIRP^7^, CHART^12^, and RAP^13^. These techniques hybridize complementary oligonucleotides to pull down a single target RNA and then identify its DNA-or protein-binding partners using next generation sequencing or mass spectrometry^13^. However, *de novo* discovery of chromatin-associated RNAs remains limited to computational predictions^1^ or association with previously known factors^14^. Alternately, nuclear fractionation allows isolation of bulk chromatin and subsequent identification of chromatin bound RNAs via sequencing, but does not provide sequence-resolved maps of RNA binding locations along the genome^15^. To overcome these limitations, we have developed ChAR-seq, a proximity ligation and sequencing method (**Figure 1A**) that both identifies chromatin-associated RNAs and maps them to genomic loci (**Figure 1B**).

**Figure 1:**
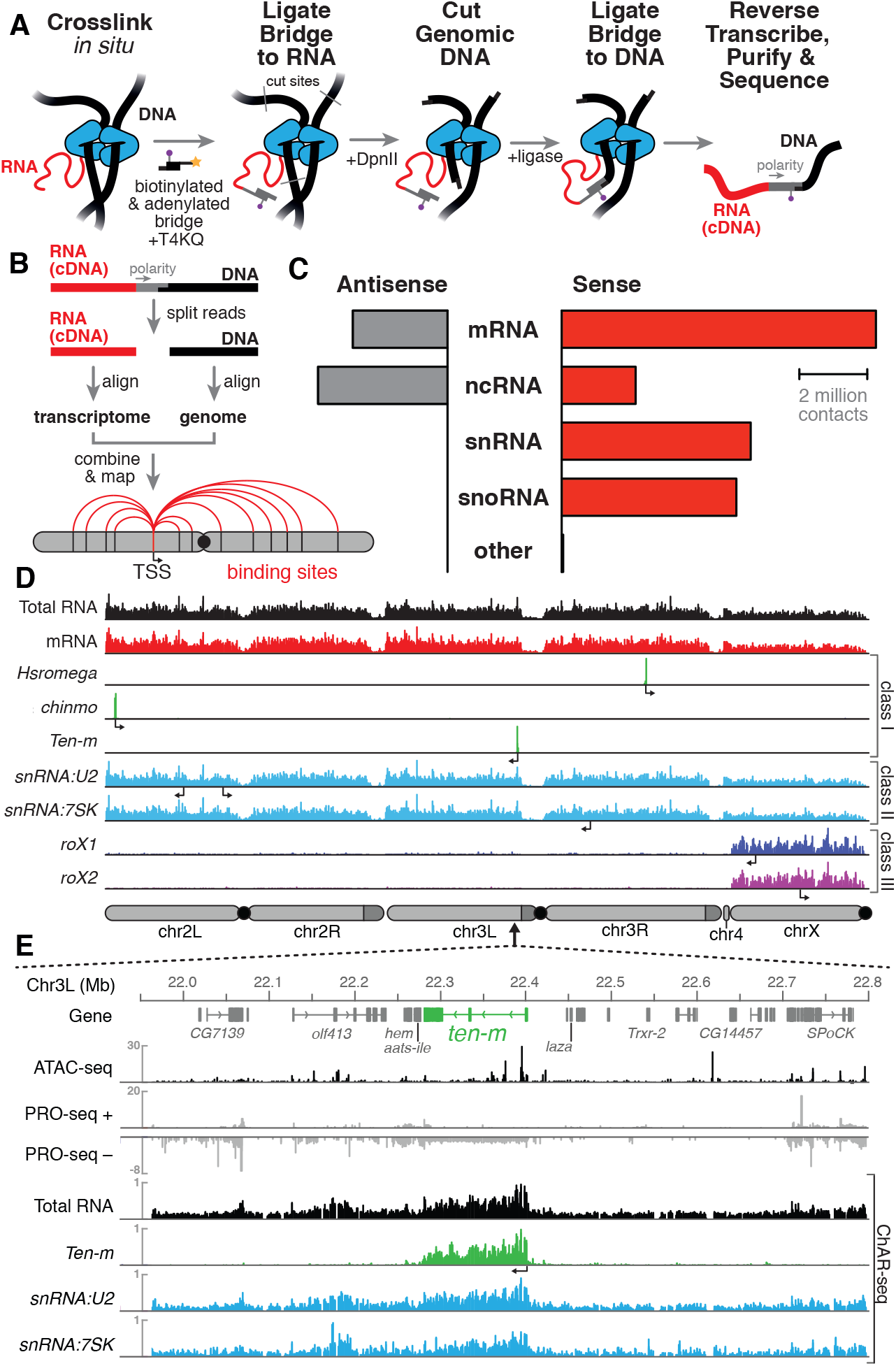
ChAR-seq uses proximity ligation of chromatin-associated RNA and deep sequencing to map RNA-DNA contacts *in situ*. **A)** Overview of the ChAR-seq method wherein RNA-DNA contacts are preserved by crosslinking, followed by *in situ* ligation of an oligonucleotide ‘bridge’ containing a 5’-adenylated ssDNA tail, biotin modification, and a DpnII-complementary overhang. The genomic DNA is then digested with DpnII and then re-ligated, capturing proximally-associated bridge molecules and RNA. The chimeric molecules are reverse-transcribed, purified and sequenced. **B)** Chimeric molecules are sequenced and the RNA and DNA ends are distinguished owing to the polarity of the bridge, which preferentially ligates to RNA via the 5’-adenylated tail and to DNA via the DpnII overhang. The RNA and DNA reads are then computationally recombined to produce contact maps for each annotated RNA in the genome. **C)** Relative abundance of chromatin-associated RNA by transcriptome classification in *Drosophila melanogaster* CME-W1-cl8+ (male) wing disc cells. **D)** Representative examples of genomewide RNA coverage plots generated for Total RNA (black), mRNA (red), *Hsromega* (green), *chinmo* (green), *ten-m* (green), *snRNA:U2* (cyan), *snRNA:7SK* (cyan), *rox1* (blue) and *roX2* (purple). Arrows show the transcription start site for each gene. **E)** Zoomed in region of shown for an 850 kilobase region of chromosome 3L (chr3L). ChAR-seq tracks for Total RNA, ten-m, snRNA:U2, and snRNA:7SK are shown in comparison with PRO-seq tracks (*Drosophila* S2^20^) and ATAC-seq (this study, CME-W1-cl8+).

## Results

We developed and performed ChAR-seq using *Drosophila melanogaster* CME-W1-cl8+ cells, a male wing disc derived cell line with a normal karyotype and well-characterized epigenome and transcriptome^16,17^. ChAR-seq is an *in situ* method^18^ for capturing genome-wide RNA-to-DNA contacts. Briefly, cells are cross-linked with formaldehyde and permeabilized, then RNA is partially fragmented and soluble RNA is washed away. The chromatin-cross-linked RNA is then ligated to an oligonucleotide duplex ’bridge’ molecule and reverse transcribed. Genomic DNA is then digested and ligated onto the other end of the oligonucleotide ’bridge’, creating a link between chromatin-associated RNA and proximal DNA. Finally, the ligated RNA is fully converted to cDNA by RNase H digestion and second strand synthesis, and the chimeric molecules are purified, processed, and sequenced.

To enable the capture and analysis of RNA-to-DNA contacts, the oligonucleotide bridge (see **Supplementary Figure 1**) was designed to have several key features: **1)** the 5’-adenylated end (5’App) enables increased ligation specificity for 3’-terminated ssRNA in the presence of truncated T4 Rnl2tr R55K K227Q ligase^19^ (**Supplementary Figure 2**), **2)** the sequence of the bridge does not exist in the yeast, fly, mouse or human genomes and encodes a defined polarity, **3)** the end of the bridge contains a restriction site for specific ligation to DNA, and **4)** the bridge is biotinylated so that it can be captured and enriched. After the bridge was ligated to RNA *in situ*, the molecules were stabilized by reverse transcription using Bst3.0 polymerase, which can traverse the DNA-RNA junction. The genomic DNA was then digested using the restriction enzyme DpnII, which produces a median fragment size of 200 bp (**Supplementary Figure 3**). The digested genomic DNA was then ligated to the bridge adaptor using T4 DNA ligase.

Upon conversion of RNA-DNA contacts to a covalent chimera, the chimeric molecules were sequenced using 152 bp single-end reads. Sequencing across the bridge junction ensures identification of the RNA and DNA portions of the chimeric molecule by reading the polarity of the bridge (**Figure 1B**). The RNA/cDNA (**Figure 1B,** *red*) and the genomic DNA side (**Figure 1B,** *black*) of each read were computationally split and aligned to the transcriptome and genome. After post-processing for unique alignments, repeat removal, and removal of blacklisted regions, each RNA molecule was mapped to the single genomic location to which it was ligated (*see* **Extended Methods** and **Supplementary Figure 4**), resulting in 24.3 million high-confidence unique mapping events for 16,817 RNA transcripts. All individual RNA-to-DNA contacts for a given transcript were then combined to produce a genome-wide association map for each individual transcript (**Figure 1B**). To ensure that ChAR-seq signal was not due to spurious bridge-to-DNA ligation, we performed a control experiment in which we added RNase A and RNase H to lysed cells before the RNA-to-bridge ligation. This RNase-treatment dramatically reduced inthe number of bridge molecules identified, demonstrating that bridge ligation is indeed RNA-dependent (**Supplementary Figure 5**).

Only the 3’-hydroxyl of each RNA is available for ligation to the bridge, thus the polarity of each RNA molecule with respect to its transcriptional direction can be determined by its orientation with respect to the polarity of the bridge. The majority (85% of total) of the RNAs captured in our assay were sense, with the largest single subtype represented by sense-stranded mRNA (32% of total), due to the capture of nascent transcripts (**Figure 1C**). Most of the chromatin-associated antisense transcripts that we identified arose from ncRNA or intronic regions. In fact, 96% of the antisense mRNAs were intronic in origin with 64% of these originating from a single 119 kb gene (*CG42339*). suggesting the presence of unannotated ncRNAs in this region. The remaining chromatin-associated RNA detected in our assay arose from non-protein coding transcription (*see* **Figure 1C**), of which 18% was small nucleolar RNA (snoRNA) and 19% was small nuclear RNA (snRNA).

ChAR-seq generated RNA-to-DNA contacts can be aggregated (**Figure 1D**, *see e.g., Total RNA*). grouped by RNA class (**Figure 1D**, *see e.g., mRNA*) or viewed individually (**Figure 1D**). Individual RNAs mapped by ChAR-seq generally fell into one of three classes. In the first class, RNAs were found around the locus from which they are transcribed (**Figure 1D**, *see, e.g., Hsromega, chinmo, ten-m*). In the second class, RNAs were found bound to chromatin in *trans,* generally distributed across most or all of the genome, often in addition to a peak around the gene body from which the RNA is transcribed (**Figure 1D**, *see e.g., snRNA:U2, snRNA:7SK*). In a third class, RNAs that are part of the dosage compensation complex (**Figure 1D**, *see roX1* and *roX2*) were enriched on and coat the X chromosome. To investigate this first class of RNAs, we compared aggregated RNA-to-DNA contacts with data from nascent transcription sequencing using PRO-seq^20^, and observed qualitative agreement between PRO-seq and ChAR-seq data sets (**Figure 1E**, *see PRO-seq* and *Total RNA).* Nevertheless, most RNA-to-DNA contacts in our dataset are associated *in trans* to genomic regions outside of the gene body from which the RNA is transcribed. For example, RNAs with strong enrichment over their own gene body, such those in class I, have on average ~20% *cis* contacts (**Supplementary Figure 6**).

ChAR-seq data can be visualized in a two-dimensional contact plot, where the genomic locus from which the RNA is transcribed is represented on the y-axis in linear genome coordinates, and the x-axis defines the genomic location where each RNA was bound. These plots provide a useful overview visualization for of the entire dataset. When we generated these contact plots for ncRNA (**Figure 2A**), mRNA (**Figure 2B**) and snRNA (**Figure 2C**), we observed strong horizontal lines that represent RNA transcripts that are transcribed from a single locus but are found distributed throughout the genome (class II), or in the special case of *roX1* and *roX2*, specifically along the X chromosome (class III). Furthermore, RNAs found at sites from which they are transcribed clustered tightly along the diagonal, a feature most pronounced for mRNAs (class I) (**Figure 2B**). Many of the RNAs we found distributed broadly across the genome are *bona fide* small nuclear RNAs (snRNAs) associated with transcription (**Figure 2C**). In fact, one of these, *snRNA:7SK,* is an abundant snRNA that functions as a scaffold for a large, transcription controlling ribonucleoprotein complex that includes p-TEFb, Hexim and LARP7, while other broadly distributed snRNAs are components of the spliceosome (e.g., *snRNA:U2*). which largely functions co-transcriptionally^21^.

**Figure 2:**
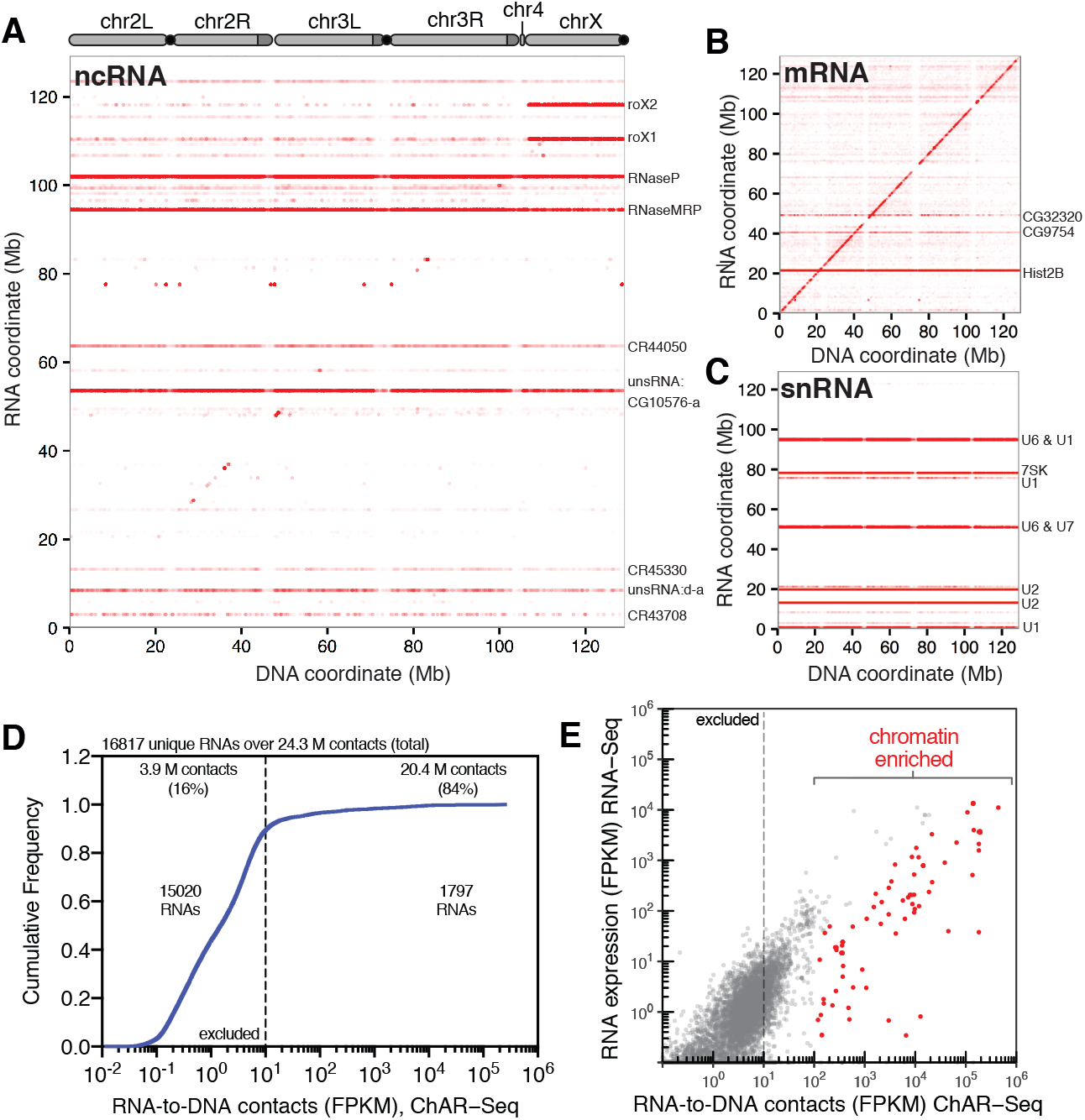
ChAR-seq is an “all to all” RNA-to-DNA proximity ligation method. **A)** Genome-wide plot of RNA to DNA contacts for non-coding RNAs. The y-axis represents the region of the genome from which a given RNA was transcribed and the x-axis represents the region of the genome where each RNA was found to be associated through proximity ligation (i.e., the binding site for each RNA). Genome-wide contact plots generated in the same way for **B)** mRNA, and **C)** snRNA. **D)** Cumulative frequency of length-normalized contacts for 16817 RNAs identified on the ‘RNA-side’ of chimeric reads. The majority (89%) of RNAs have fewer than 10 fragments per kilobase per million reads (FPKM) in our dataset and were not further analyzed owing to low coverage. The remainder of the 1797 RNAs account for 20.4 (84%) of the total RNA-to-DNA contacts. **E)** Scatter plot of length normalized chromatin-contacts versus total expression for each RNA.

To identify RNAs highly enriched for substantial chromatin interactions, we plotted the normalized cumulative distribution of the number of contacts observed for each gene (**Figure 2D**). The majority of the RNAs in our dataset (15,020 out of 16817, 89%) had fewer than 10 FPKM contacts (**Figure 2D**), and were excluded from further analysis. The remainder of the 1797 RNAs (11%) accounted for 84% (20.4 million) of all chromatin contacts in our data set. To estimate the contribution of total RNA abundance to this interaction signal, we performed RNA-seq to for the CME-W1-cl8+ cell line and compared RNA expression levels with RNA-to-DNA contacts identified by ChAR-seq (**Figure 2E**). Unsurprisingly, we observed a correlation between RNA expression and chromatin-RNA contacts; however, a cluster of RNAs clearly generated more chromatin interactions that would be expected from the overall expression levels (**Figure 2E**). Using both the number of contacts and the fold-enrichment over RNA expression, we identified 73 RNAs that had more than 100 FPKM contacts and were enriched more than four-fold above expectation, though many were enriched by 2-5 orders of magnitude (**Figure 2E**, red symbols; **Supplementary Figure 7-8**).

We developed ChAR-seq using the male WME-cl8+ line, reasoning that the ncRNAs *roX1* and *roX2* would serve as an internal positive control. Both *roX1* and *roX2* are part of the MSL2 complex, which binds across the X-chromosome in male flies to recruit chromatin-modifiers that increase transcriptional output (**Figure 3A**)^22^. Indeed, ChAR-seq data showed *roX1* and *roX2* to be 7.6-fold (p-value < 10^−10^) and 8.1-fold (p-value < 10^−10^) enriched for interactions on the X chromosome, respectively (**Figure 3B,C**). In contrast, female flies express *Sex lethal* (*Sxl*). which binds to *msl2* mRNA to prevent its translation, blocking assembly of the MSL2 complex^22^. Importantly, *roX1* and *roX2* require MSL2 for X-chromosome specific localization^22^, therefore female cells should lack detectable spreading of these ncRNAs along the X-chromosome. When we performed ChAR-seq in a female *Drosophila melanogaster* cell line, Kc167, we did not detect any significant *roX2* localization on the X chromosome (**Figure 3D**), but observed excellent agreement in interaction signal from other RNAs across both cell lines (**Supplementary Figure 9, Figure 3C** *male, CME-W1-cl8+* and **Figure 3D**, *female, Kc167, see e.g., snRNA: 7SK* and *Hsromega*).

**Figure 3:**
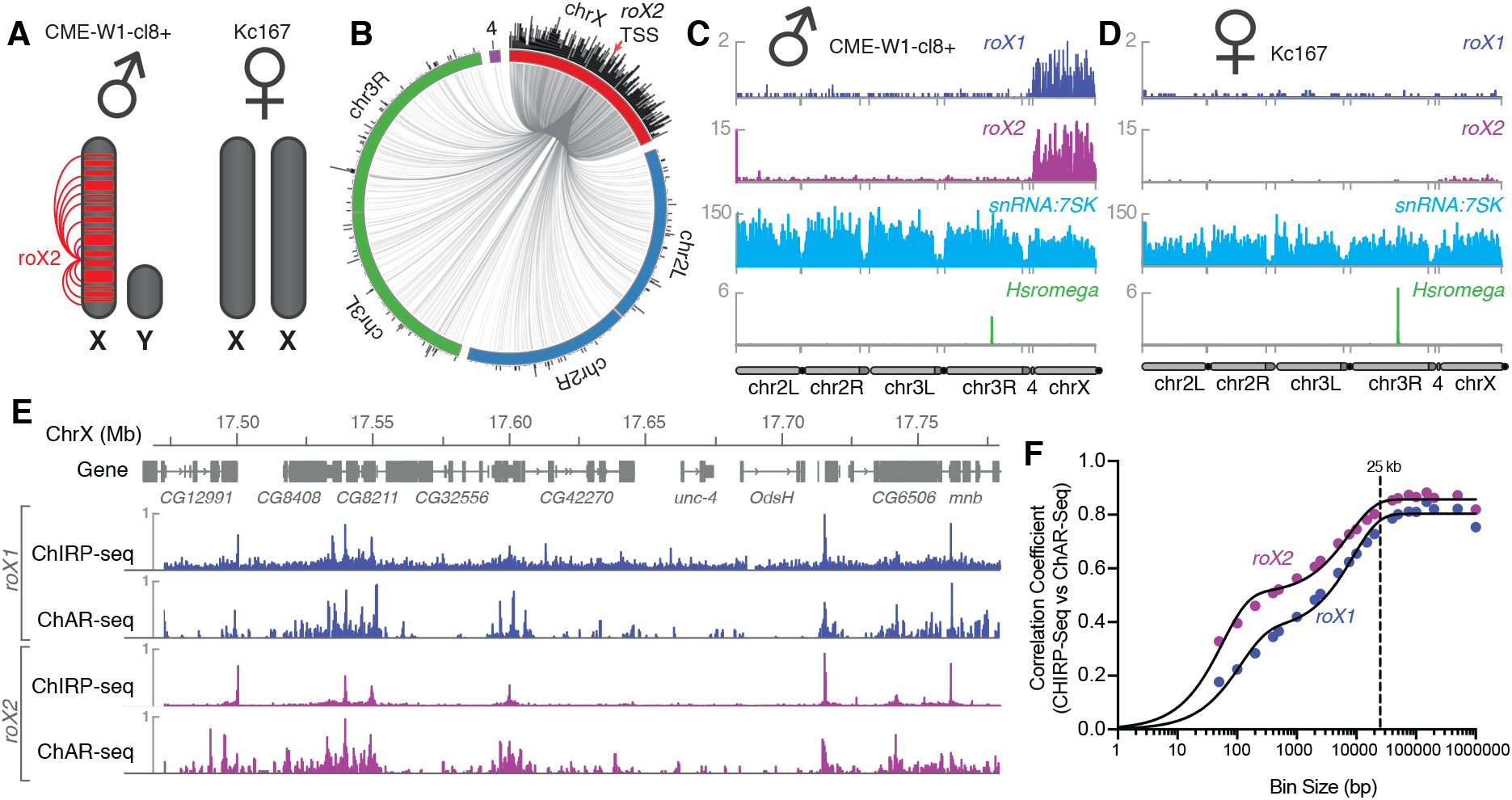
Mapping *roX1* and *roX2* of the X chromosome dosage compensation complex. **A)** Illustration of the *roX1/roX2* spreading across the solitary X chromosome in male flies (CME-W1-cl8+ cell line). In contrast, the female-derived Kc167 cell line expresses significantly lower levels of the MSL2 complex, which mediates the association of *roX1* and *roX2,* which therefore do not coat either of the two X-chromosomes in females. **B)** Circos plot showing *roX2* spreading from its site of transcription (red arrow) and binding with high density along the X-chromosome but low density binding throughout the genome. **C)** Coverage plots of *roX1* (blue), *roX2* (purple), *snRNA:7SK* (cyan) and *Hsromega* (green) in male CME-W1-cl8+ cells. **D)** Complementary coverage plots generated from female Kc167 cells. **E)** Comparison of ChAR-seq (*this work*) to an alternative RNA-to-chromatin mapping method called ChIRP-seq (data from reference 23). Tracks for *roX1* (upper, blue) and *roX2* (lower, purple) were generated from 32308 and 87453 contacts, respectively, from a ChAR-seq dataset containing a total of 24.7 million contacts. For comparison, the *roX1* and *roX2* tracks derived from ChIRP-seq each represent greater that 20 million reads. **F)** Correlation coefficients were calculated for *roX1* and *roX2* coverage tracks generated using ChIRP-seq and ChAR-seq and plot relative to increasing bin size to estimate the resolution of the assay.

High-resolution maps of *roX1* and *roX2* localization have previously been generated using ChIRP-Seq, which hybridizes probes against a known RNA and pulls down the associated chromatin for sequencing^7,23^. Comparing ChIRP-seq to ChAR-seq for both *roX1* and *roX2* (**Figure 3E**), we found that DNA contact locations were in surprisingly good agreement despite the fact that ChAR-seq reads are spread across all RNAs while ChIRP-seq reads map the specific RNA target, resulting in a large disparity in the effective sequencing depth between the methods. In ChIRP-seq, virtually all of the signal is attributable to interactions between chromatin and the target RNA. In contrast, ChAR-seq captures *all* RNA and DNA contacts, so that any given target RNA will comprise a subset of the total RNA-chromatin contacts in the dataset. In the case of *roX1* and *roX2,* we observed 32,308 and 87,453 contacts, representing 0.1% and 0.36% of the ChAR-seq dataset. In contrast, the ChIRP-seq datasets plotted in **Figure 3E** represent ~24M and ~21M reads for *roX1* and *roX2*, respectively. This indicates that ChAR-seq can identify RNA peaks along chromatin with high sensitivity for a given RNA.

The resolution with which we can measure the localization of an RNA to a given genomic site constrains our ability to assess its potential modes of action. To measure the accuracy of ChAR-seq measurements of RNA interaction with DNA, we used the ChIRP-seq data set to calculate the base-pair resolution of the method. We expected this resolution to be bounded—in part—by the local DpnII cut frequency (**Supplementary Figure 3**) and the number of contacts for any given RNA. We divided the X chromosome into evenly sized bins and calculated correlation coefficients between ChIRP-seq and ChAR-seq datasets at increasing bin sizes for both *roX1* and *roX2* (**Figure 3F**). Using this method, we noted a bi-phasic increase of the correlation coefficient, corresponding to a minor plateau around 200 bp and a major plateau at ~25 kbp. The minor plateau is likely due to the DpnII distribution bias in the ChAR-seq tracks, while the major plateau is an estimate of the resolution of our assay, which is on the order of other proximity-ligation sequencing assays like Hi-C^24^.

To test if we could identify the functional roles for our most highly enriched RNAs, we clustered the snRNA class of RNAs based on their genomic contacts. These snRNAs collectively comprised 23% of all the RNA-to-DNA contacts in our dataset (**Figure 4A**) and are a substantial component of the spliceosome, a multi-megadalton ribonucleoprotein complex that catalyzes pre-mRNA splicing^25,26^. The composition and conformation of the spliceosome is highly dynamic, though two dominant species exist in eukaryotes: the major spliceosome comprised of U1, U2, U4, U5 and U6 snRNAs, and the minor spliceosome comprised of U4:atac, U6:atac, U5, U11, and U12^26^. Many members of this class of snRNAs have highly similar gene duplication variants in the *Drosophila* genome. We therefore first calculated the base sequence similarity of these variants to one another and aggregated signals that were tightly clustered (**Supplementary Figure 10**). When we then correlated genome-wide binding signal within this class, we found that the distribution patterns of the major spliceosome snRNAs U1, U2, U4, U5, U6 clustered together along with *snRNA:7SK* (**Figure 4B**), which is part of the p-TEFb complex that relieves pausing of RNA Polymerase II at promoters^27^ and may participate in the release of paused polymerase during RNA splicing ^28^. The components of the minor spliceosome did not cluster together, likely due to their low abundance^26^ and consequently low representation in our dataset.

**Figure 4:**
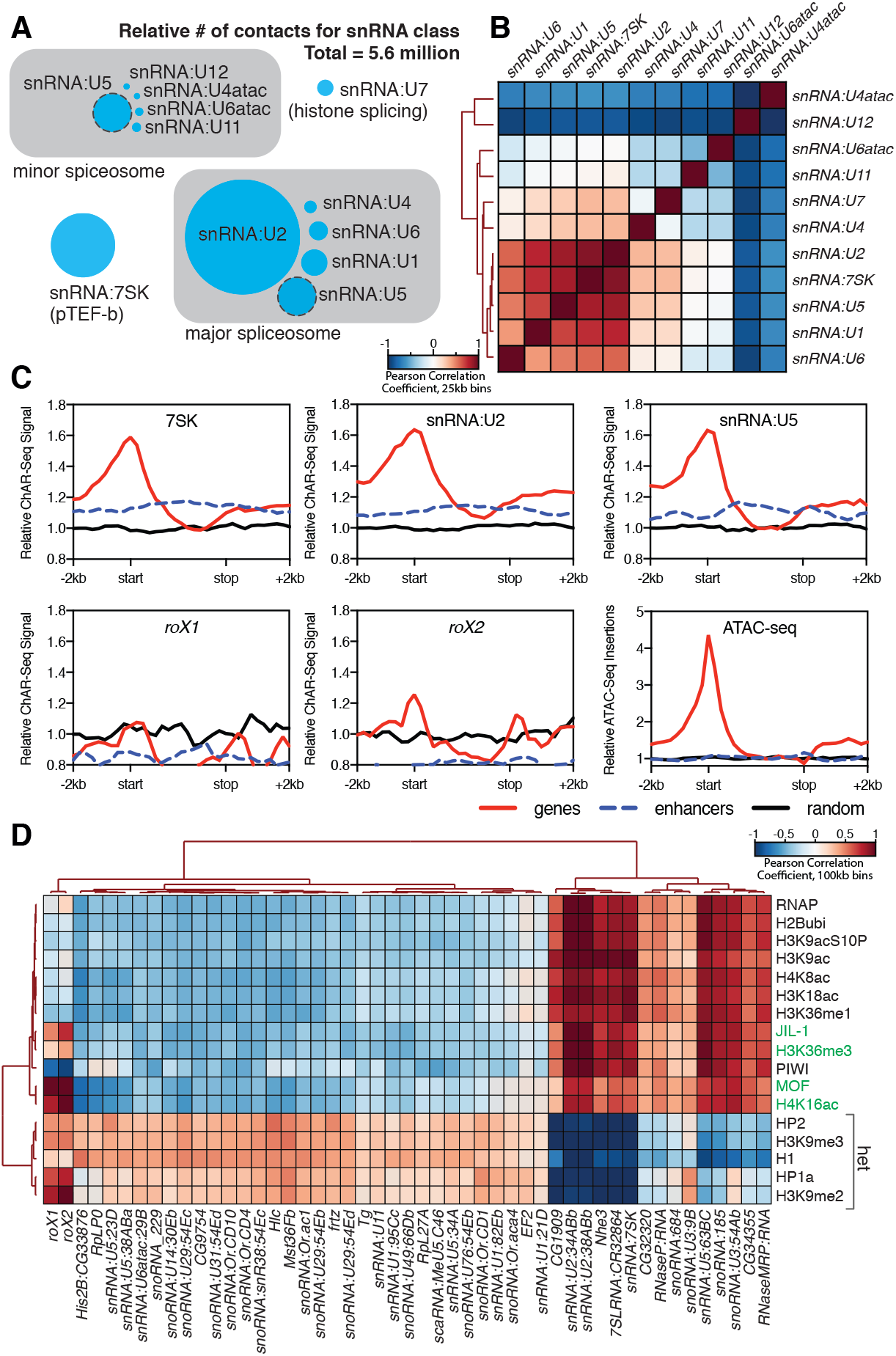
Correlation of chromatin-associated RNAs with genome features. **A)** Relative abundance of snRNAs identified by ChAR-seq. The size of the circles is proportional to the abundance of the snRNAs found by ChAR-seq. RNA components of the major and minor spliceosome are bounded by the gray boxes. **B)** Cluster analysis of the pairwise correlation between genome-wide tracks of snRNAs. **C)** Metaanalysis plots aggregating the signal of *snRNA:7SK, snRNA:U2, snRNA:U5, roX1, roX2* and ATAC-seq over gene bodies (red), putative enhancers (blue dashed line) and random regions (black). **D)** Hierarchical clustering based on pairwise Pearson correlation between representative ChAR-seq RNA-to-DNA contact coverage tracks (black) and modENCODE datasets available for the WME-C1-cl8+ cell line. Notable associations for the dosage compensation complex (green) and heterochromatin (“het”) are indicated in the right margin.

We next reasoned that spliceosome RNAs—as part of the co-transcriptional RNA processing machinery—should also be enriched in gene bodies. We therefore aggregated spliceosomal RNA signals over gene bodies (**Figure 4C,** *red lines*). putative enhancers^29^ (**Figure 4C,** *blue dashed lines*) and a random distribution of genomic bins of similar size (**Figure 4C,** *black lines*). We observed an enrichment of snRNAs (7SK, U2 and U6), but not *roX1* or *roX2,* over gene bodies (**Figure 4C**) with a broad peak around transcription start sites, in good agreement with ChIRP data for 7SK in mice^30^.

In contrast to the small number of well-defined and well-characterized snRNAs involved in splicing, there are more than 200 snoRNAs in flies^31^ that are significantly divergent in sequence and, surprisingly, were highly represented in our dataset (**Supplementary Figure 7-8** and **Figure 1C**). Most of these snoRNAs have either unknown function or are computationally identified and indirectly implicated in the maturation and modification of ribosomal rRNA^31^.

To determine if our enriched chromatin-associated RNAs, in particular snRNAs and snoRNAs, might localize to euchromatic or heterochromatic states or with specific transcription factors, we cross-correlated our ChAR-seq signal against modENCODE datasets available for the CME-W1-cl8+ cell line. To normalize the signals for comparison, we first calculated the expected contacts per 2 kb bin for each RNA under a uniform distribution, based on the total number of genome-wide contacts for each RNA and the number of DpnII sites per bin. This null model was then used to calculate the log2 ratio of the observed to the expected contacts per bin for each RNA, which was then transformed into a z-score ((x−μ)/σ) based on the whole-genome mean (μ) and standard deviation (σ). Similarly, we re-binned the modENCODE tracks, removed bins that did not contain a DpnII site, and transformed the log2 mean-shift values to a z-score. We then calculated the pair-wise Pearson correlation coefficients between each signal track, and then clustered the data (**Figure 4D**). We observed discrete clustering of *roX1* and *roX2* with known dosage compensation complex factors, MOF, the histone modifications H4K16ac and H3K36me3^32^, and JIL-1 kinase^33^, validating this analytical approach (**Figure 4D**). Beyond this sub-cluster of dosage compensation factors, the remainder of the chromatin-associated RNAs fell into two distinct and anti-correlated categories: those associated with active chromatin and transcription (e.g., RNAPII, H4K8ac, H3K18ac) or heterochromatin (e.g., HP2, H3K9me3, HP1a) (**Figure 4D**). In particular, we note that *snRNA:U2* and *snRNA:7SK* cluster tightly with the transcription-associated chromatin marks, while many of the snoRNAs and minor spliceosome snRNAs that we identified are associated with heterochromatin, likely due to co-localization of heterochromatin factors to the nucleolus. Interestingly, *snRNA:U5,* a component of both the major and minor spliceosome, has variants that clearly cluster with either transcriptionally active chromatin (63BC) or heterochromatin (23D, 38ABa, and 34A). Previous work has shown that the *snRNA:U5:38Aba* variant (**Figure 4D**, heterochromatin cluster) exhibits a unique tissue-specific expression profile with the greatest abundance in neural tissue, which led the authors to propose isoform-dependent functions in alternative splicing^34^. The differential clustering that we observe for *snRNA:U5,* and indeed between major and minor spliceosome snRNAs, between euchromatin and heterochromatin might reflect such isoform-specific functions of the spliceosome in different chromatin states.

## Discussion

ChAR-seq maps the chromosomal binding sites of all chromatin-associated RNAs, independent of whether they are associated as nascent transcripts or bound as part of ribonucleoprotein complexes (RNPs). In this way, ChAR-seq can be thought of as a massively parallelized *de novo* RNA mapping assay capable of generating hundreds to thousands of RNA-binding maps. ChAR-seq also detects multiple classes of chromatin-associated RNAs. We validated ChAR-seq using chromosome-specific ncRNAs *roX1* and *roX2* associated with dosage compensation. The comparison between ChAR-seq and ChIRP-seq, which vary dramatically in the sequencing depth needed to analyze a specific RNA, highlights the utility of ChAR-seq as a de novo chromatin-associated RNA discovery tool. ChAR-seq also maps nascent RNAs found at the loci from which they are transcribed. ChAR-seq is similar to a recently published method^38^, but has two key distinctions. First, proximity ligations are performed *in situ* in intact nuclei, which reduces nonspecific interactions^18^. Second, ChAR-seq uses long single-end reads to sequence across the entire junction of the ‘bridge’, ensuring that RNA-to-DNA contacts are mapped with high confidence and reporting on the polarity of the bridge-ligated RNA.

We used ChAR-seq to discover and map several dozen ncRNAs that are pervasively bound across the genome. Many of these ncRNAs are components of ribonucleoprotein complexes associated with transcription elongation (*snRNA:7SK*). splicing (*snRNA:U2,* etc) and RNA processing (*snRNAs, snoRNAs* and *scaRNAs*). Interestingly, more than half of the chromatin-associated RNAs identified based on our enrichment criteria are snoRNAs, most of which—but not all—correlate with heterochromatin. Generally, snoRNA ribonucleoproteins (snoRNPs) use intermolecular base pairing to direct chemical modification of the 2’-hydroxyl groups or the isomerization of uridines to pseudouridine^3^ and snoRNAs are known abundant components of chromatin in both flies^35^ and in mice^36^. Despite their abundance and the their known role in RNA modification, we do not yet understand the functions of these modifications, or the implication of the abundance of snoRNAs and scaRNAs in cells or associated with chromatin^3^. Finally, we demonstrate that ChAR-seq can be used with orthogonal genome-wide datasets to identify and classify RNAs that are associated with specific chromatin states (e.g., euchromatin vs heterochromatin), which we expect will be particularly useful in higher organisms that use lncRNAs such as *HOTAIR, HOTTIP* and *BRAVEHEART* as scaffolds for ribonucleoproteins that regulate facultative heterochromatin.

We anticipate that ChAR-seq will be a powerful new high throughput discovery platform capable of simultaneously identifying new chromatin-associated RNAs and mapping their chromatin binding sites (and associated epigenetic chromatin states), all of which will be particularly useful in comparing ‘epigenomic’ changes that coincide with cellular differentiation and/or tumorigenesis.

## Acknowledgements

This project was funded by a Stanford Center for Systems Biology (NIH P50 GM107615) Seed Grant to JCB, DJ, VIR & WLJ. JCB and DJ were also supported by NIH Ruth Kirchstein National Research Service Awards (F32GM116338 to JCB) and (F32GM108295 to DJ). JCB was also supported by the Stanford School of Medicine Dean’s Fellowship. VIR was supported by the Walter V. and Idun Berry Fellowship. NAT was supported by the Stanford Genetics Training Program (5T32HG000044-19). OKS was supported by the Molecular Pharmacology Training Grant (NIH T32-GM113854-02). WLJ was supported by a NIH T32 Training Fellowship (GM007276) and the National Science Foundation Graduate Research Fellowship (DGE-114747). We acknowledge support from NIH RO1 HD085135 to JMS and AFS, HHMI-Simons Faculty Scholar Award to JMS, NIH grants P50HG00773501 and R21HG007726 to WJG, and R01GM106005 to AFS. We would like to thank Julia Salzman, Kyle Eagen, and members of the Straight, Greenleaf and Skotheim labs for thoughtful discussions. We thank Lucy O’Brien for sharing equipment. We acknowledge the Drosophila Genomics Resource Center (NIH grant 2P40OD010949-10A1) for providing cell lines and the Stanford Functional Genomics Facility for providing sequencing services.

## Author contributions

JCB, VIR, WLJ and DJ conceived of the idea and planned experiments. JCB, NAT and OKS prepared all ChAR-seq libraries. DJ prepared ATAC-seq libraries. JCB, NAT, VIR, DJ and OKS processed and analyzed data. AFS, WJG, and JMS provided advice and material support. All authors discussed and interpreted the results and contributed to the writing and editing of the manuscript.

## Method Summary

*Drosophila melanogaster* CME-W1-cl8+ cells (Drosophila Genome Resource Center) were grown in T-75 flasks at 27°C in Shields & Sang M3 media supplemented with 5 μ g/mL insulin, 2% FBS, 2% fly extract and 100 μg/mL Pen-Strep^16^. Approximately 100-400 million cells were harvested for each library by centrifugation at 2000 × g for 2-4 minutes, resuspended in fresh media plus 1% formaldehyde and fixed for 10 minutes at room temperature. Fixation was quenched by adding 0.2 M glycine and mixing for 5 minutes at room temperature. Cells were centrifuged at 2000 × g for 2-4 minutes, resuspended in 1 mL of PBS, and centrifuged again at 2000 × g for 2 minutes. The supernatant was aspirated and discarded, and the cell pellet was flash frozen in liquid nitrogen and stored at −80C until needed. Cells were thawed in lysis buffer and the cross-linked nuclei and cellular material were isolated by centrifugation for the *in situ* ligation protocol (see *Extended Methods for details).*

Briefly, RNA was lightly and partially chemically fragmented by heating in the presence of magnesium. The pellet was isolated and washed, and RNA ends were ligated using truncated T4 Rnl2tr R55K K227Q ligase (hereafter referred to as trT4KQ RNA ligase) to an oligonucleotide ’bridge’ molecule containing a 5’-adenylated ssDNA overhang. The RNA ligase was inactivated, the pellet was washed and the RNA strand was stabilized by first strand synthesis of the RNA through extension of the bridge by Bst 3.0 polymerase. The polymerase was inactivated and the pellet was washed. Genomic DNA was then digested with DpnII, followed by ligation of the DpnII digested genomic DNA to the opposite end of the oligonucleotide ’bridge’. Second strand synthesis was then performed using RNaseH and DNA Polymerase I to complete cDNA synthesis of the RNA-encoded side of the new, chimeric molecules. The sample was then deproteinized and crosslinks were reversed by heating overnight in SDS and proteinase K. DNA was then ethanol precipitated and sheared to ~200 bp fragments using a Covaris focused ultra-sonicator. DNA fragments containing the biotinylated bridge were then purified using magnetic streptavidin-coated beads. DNA ends were repaired using the NEBNext End Repair and dA tailing module, and ligated to NEBNext hairpin adaptors for Illumina sequencing. The adaptor hairpin was cleaved using USER, and DNA fragments were amplified by ∼8-12 rounds of PCR with NEBNext Indexing Primers for Illumina (TruSeq compatible). The partially amplified library was then purified using AmPure XP beads to remove adaptor dimers, and the optimum number of additional PCR cycles was determined by qPCR to achieve approximately 30% saturation. Library amplification was then completed by the additional rounds of PCR, and the library was purified and size selected to a target range of 100-500 bps using AmPure XP beads. The size distribution of the library was checked by capillary electrophoresis using an Agilent Bioanalyzer, and quantified using qPCR against a phiX Illumina library standard curve. Libraries were sequenced using the Illumina MiSeq platform for quality control, and subsequently sequenced on the Illumina NextSeq platform (Stanford Functional Genomics Facility) using single-end 152 bp reads. Data was processed and analyzed using a custom pipeline (see *Extended Methods for details*).

## References

1 Guttman, M. & Rinn, J. L. Modular regulatory principles of large non-coding RNAs. Nature 482, 339-346, doi:10.1038/nature 10887 (2012).

2 Meller, V. H., Joshi, S. S. & Deshpande, N. Modulation of Chromatin by Noncoding RNA. Annu Rev Genet 49, 673-695 (2015).

3 Cech, T. R. & Steitz, J. A. The noncoding RNA revolution-trashing old rules to forge new ones. Cell 157, 77-94 (2014).

4 Conrad, T. & Akhtar, A. Dosage compensation in Drosophila melanogaster: epigenetic fine-tuning of chromosome-wide transcription. Nat Rev Genet 13, 123-134, doi:10.1038/nrg3124 (2012).

5 Augui, S., Nora, E. P. & Heard, E. Regulation of X-chromosome inactivation by the X-inactivation centre. Nat Rev Genet 12, 429-442, doi:10.1038/nrg2987 (2011).

6 Rinn, J. L. et al. Functional demarcation of active and silent chromatin domains in human HOX loci by noncoding RNAs. Cell 129, 1311-1323, doi:10.1016/j.cell.2007.05.022 (2007).

7 Chu, C., Qu, K., Zhong, F. L., Artandi, S. E. & Chang, H. Y. Genomic maps of long noncoding RNA occupancy reveal principles of RNA-chromatin interactions. Mol Cell 44, 667-678, doi: 10.1016/j.molcel.2011.08.027 (2011).

8 Wang, K. C. et al. A long noncoding RNA maintains active chromatin to coordinate homeotic gene expression. Nature 472, 120-124, doi:10.1038/nature09819 (2011).

9 Sigova, A. A. et al. Transcription factor trapping by RNA in gene regulatory elements. Science 350, 978-981, doi:10.1126/science.aad3346 (2015).

10 Bunting, S. F. et al. 53BP1 inhibits homologous recombination in Brca1-deficient cells by blocking resection of DNA breaks. Cell 141, 243-254, doi:10.1016/j.cell.2010.03.012 (2010).

11 Hall, L. E., Mitchell, S. E. & O’Neill, R. J. Pericentric and centromeric transcription: a perfect balance required. Chromosome Res 20, 535-546, doi:10.1007/s10577-012-9297-9 (2012).

12 Engreitz, J. M. et al. The Xist lncRNA exploits three-dimensional genome architecture to spread across the X chromosome. Science 341, 1237973, doi:10.1126/science.1237973 (2013).

13 Simon, M. D. et al. The genomic binding sites of a noncoding RNA. Proc Natl Acad Sci U S A 108, 20497-20502, doi:10.1073/pnas.1113536108 (2011).

14 Khalil, A. M. et al. Many human large intergenic noncoding RNAs associate with chromatin-modifying complexes and affect gene expression. Proc Natl Acad Sci U S A 106, 11667-11672, doi: 10.1073/pnas.0904715106 (2009).

15 Werner, M. S. & Ruthenburg, A. J. Nuclear Fractionation Reveals Thousands of Chromatin-Tethered Noncoding RNAs Adjacent to Active Genes. Cell Rep 12, 1089-1098, doi:10.1016/j.celrep.2015.07.033 (2015).

16 Cherbas, L. et al. The transcriptional diversity of 25 Drosophila cell lines. Genome Res 21, 301-314, doi:10.1101/gr. 112961.110 (2011).

17 modENCODE Consortium, Identification of functional elements and regulatory circuits by Drosophila modENCODE. Science 330, 1787-1797, doi:10.1126/science.1198374 (2010).

18 Rao, S. S. et al. A 3D map of the human genome at kilobase resolution reveals principles of chromatin looping. Cell 159, 1665-1680, doi:10.1016/j.cell.2014.11.021 (2014).

19 Viollet, S., Fuchs, R. T., Munafo, D. B., Zhuang, F. & Robb, G. B. T4 RNA ligase 2 truncated active site mutants: improved tools for RNA analysis. BMC biotechnology 11, 72, doi:10.1186/1472-6750-11-72 (2011).

20 Kwak, H., Fuda, N. J., Core, L. J. & Lis, J. T. Precise maps of RNA polymerase reveal how promoters direct initiation and pausing. Science 339, 950-953, doi:10.1126/science. 1229386 (2013).

21 Perales, R. & Bentley, D. “Cotranscriptionality”: the transcription elongation complex as a nexus for nuclear transactions. Mol Cell 36, 178-191, doi:10.1016/j.molcel.2009.09.018 (2009).

22 Lucchesi, J. C. & Kuroda, M. I. Dosage compensation in Drosophila. Cold Spring Harb Perspect Biol 7, doi:10.1101/cshperspect.a019398 (2015).

23 Quinn, J. J. et al. Revealing long noncoding RNA architecture and functions using domain-specific chromatin isolation by RNA purification. Nat Biotechnol 32, 933-940, doi:10.1038/nbt.2943 (2014).

24 van Berkum, N. L. et al. Hi-C: a method to study the three-dimensional architecture of genomes. J Vis Exp, doi:10.3791/1869(2010).

25 Zhou, Z., Licklider, L. J., Gygi, S. P. & Reed, R. Comprehensive proteomic analysis of the human spliceosome. Nature 419, 182-185, doi:10.1038/nature01031 (2002).

26 Will, C. L. & Luhrmann, R. Spliceosome structure and function. Cold Spring Harb Perspect Biol 3, doi:10.1101/cshperspect.a003707 (2011).

27 Kwak, H. & Lis, J. T. Control of transcriptional elongation. Annu Rev Genet 47, 483-508, doi:10.1146/annurev-genet-110711-155440 (2013).

28 Barboric, M. et al. 7SK snRNP/P-TEFb couples transcription elongation with alternative splicing and is essential for vertebrate development. Proc Natl Acad Sci U S A 106, 7798-7803, doi:10.1073/pnas.0903188106 (2009).

29 Kvon, E. Z. et al. Genome-scale functional characterization of Drosophila developmental enhancers in vivo. Nature 512, 91-95, doi:10.1038/nature13395 (2014).

30 Flynn, R. A. et al. 7SK-BAF axis controls pervasive transcription at enhancers. Nat Struct Mol Biol 23, 231-238 (2016).

31 Huang, Z. P. et al. Genome-wide analyses of two families of snoRNA genes from Drosophila melanogaster, demonstrating the extensive utilization of introns for coding of snoRNAs. RNA 11, 1303-1316, doi:10.1261/rna.2380905 (2005).

32 Bell, O. et al. Transcription-coupled methylation of histone H3 at lysine 36 regulates dosage compensation by enhancing recruitment of the MSL complex in Drosophila melanogaster. Mol Cell Biol 28, 3401-3409, doi:10.1128/MCB.00006-08 (2008).

33 Bai, X., Alekseyenko, A. A. & Kuroda, M. I. Sequence-specific targeting of MSL complex regulates transcription of the roX RNA genes. EMBO J 23, 2853-2861, doi:10.1038/sj.emboj.7600299 (2004).

34 Chen, L. et al. Identification and analysis of U5 snRNA variants in Drosophila. RNA 11, 1473-1477, doi:10.1261/rna.2141505 (2005).

35 Schubert, T. et al. Df31 protein and snoRNAs maintain accessible higher-order structures of chromatin. Mol Cell 48, 434-444 (2012).

36 Meng, Y. et al. The non-coding RNA composition of the mitotic chromosome by 5’-tag sequencing. Nucleic Acids Res 44, 4934-4946, doi:10.1093/nar/gkw195 (2016).

37 Booth, D. G. et al. 3D-CLEM Reveals that a Major Portion of Mitotic Chromosomes Is Not Chromatin. Mol Cell 64, 790-802, doi:10.1016/j.molcel.2016.10.009 (2016).

38 Sridhar, B. et al. Systematic Mapping of RNA-Chromatin Interactions In Vivo. Curr Biol 27, 602-609, doi:10.1016/j.cub.2017.01.011 (2017).

39 Petersen, K. R., Streett, D. A., Gerritsen, A. T., Hunter, S. S. & Settles, M. L. in Proceedings of the 6th ACM Conference on Bioinformatics, Computational Biology and Health Informatics 491-492 (ACM, Atlanta, Georgia, 2015).

40 Bolger, A. M., Lohse, M. & Usadel, B. Trimmomatic: a flexible trimmer for Illumina sequence data. Bioinformatics 30, 2114-2120, doi:10.1093/bioinformatics/btu170 (2014).

41 Langmead, B. & Salzberg, S. L. Fast gapped-read alignment with Bowtie 2. Nat Methods 9, 357-359, doi:10.1038/nmeth. 1923 (2012).

42 Li, H. et al. The Sequence Alignment/Map format and SAMtools. Bioinformatics 25, 2078-2079, doi:10.1093/bioinformatics/btp352 (2009).

43 Quinlan, A. R. & Hall, I. M. BEDTools: a flexible suite of utilities for comparing genomic features. Bioinformatics 26, 841-842, doi:10.1093/bioinformatics/btq033 (2010).

44 Zhang, Y. et al. Model-based analysis of ChIP-Seq (MACS). Genome Biol 9, R137, doi:10.1186/gb-2008-9-9-r137 (2008).

45 Krzywinski, M. et al. Circos: an information aesthetic for comparative genomics. Genome Res 19, 1639-1645, doi:10.1101/gr.092759.109 (2009).

46 Ramirez, F. et al. deepTools2: a next generation web server for deep-sequencing data analysis. Nucleic Acids Res 44, W160-165, doi:10.1093/nar/gkw257 (2016).

47 Heinz, S. et al. Simple combinations of lineage-determining transcription factors prime cis-regulatory elements required for macrophage and B cell identities. Mol Cell 38, 576-589, doi: 10.1016/j.molcel.2010.05.004 (2010).

48 Kearse, M. et al. Geneious Basic: an integrated and extendable desktop software platform for the organization and analysis of sequence data. Bioinformatics 28, 1647-1649, doi:10.1093/bioinformatics/bts199 (2012).

49 ENCODE Consortium, An integrated encyclopedia of DNA elements in the human genome. Nature 489, 57-74, doi:10.1038/nature11247 (2012).

50 Buenrostro, J. D., Giresi, P. G., Zaba, L. C., Chang, H. Y. & Greenleaf, W. J. Transposition of native chromatin for fast and sensitive epigenomic profiling of open chromatin, DNA-binding proteins and nucleosome position. Nat Methods 10, 1213-1218, doi:10.1038/nmeth.2688 (2013).

